# Prevalence of Key Spike Protein Mutations and Their Limited Effect on COVID-19 Clinical Manifestations in Sylhet, Bangladesh

**DOI:** 10.1101/2024.10.25.620187

**Authors:** Shahina Akter, Zebunnesa Zeba, Md. Anwar Hossain, Michael Arthur Ofori, Muhammad Shahab, Gabriel Vinícius Rolim Silva, Jonas Ivan Nobre Oliveira, Shopnil Akash, Tanjina Akhtar Banu, Mousona Islam, Murshed Hasan Sarkar, Barna Goswami, Sanjana Fatema Chowdhury, Md. Ahashan Habib, Md. Salim Khan

**Author notes:** Corresponding author Email: Dr. Shahina Akter.

## Abstract

SARS-CoV-2 is the virus responsible for the COVID-19 pandemic, which has spread rapidly around the world and had a significant impact on public health and the economy worldwide. This study investigated the correlation between SARS-CoV-2 spike protein mutations, clinical outcomes and patient demographics in the Sylhet region of Bangladesh. We looked at the full genome sequences of 37 SARS-CoV-2 samples that were collected between January and June 2020. Specifically, we looked at five major spike protein mutations: D614G, A570D, D1118H, A222V, and P681R. The D614G mutation was the most prevalent (94.6%), followed by A570D and D1118H (both 32.4%), A222V (29.7%), and P681R (13.5%). Despite their high prevalence, we found no statistically significant associations between these mutations and clinical outcomes or demographic variables, except for possible trends for the P681R mutation. We found that age played a decisive role in recovery from COVID-19, with older patients exhibiting slower recovery rates. In terms of predictors of outcome, gender differences were observed: clinical symptoms and viral genetic mutations were more influential for men, while age and disease progression were more important for women. Common nucleotide substitutions (A23403G, C3037T, and C14408T) associated with European strains were identified, suggesting possible routes of transmission. This study contributes to our understanding of the genetics, clinical manifestations and epidemiology of SARS-CoV-2 in the Sylhet region and emphasizes the need for continuous genomic surveillance and adaptive public health strategies.

## Introduction

Since its emergence in late 2019, the novel coronavirus SARS-CoV-2, responsible for the COVID-19 pandemic, has triggered an unprecedented global health crisis (Lai et al., 2020; Akter et al. 2022). The virus’s rapid human-to-human transmission has led to widespread morbidity and mortality, severely impacting global economies and healthcare systems (Zhu et al., 2020; Sarkar et al. 2023). As the pandemic has progressed, numerous mutations have emerged in the SARS-CoV-2 genome, particularly in the spike protein, which may influence viral transmissibility, pathogenicity, and immune evasion (Lauring & Hodcroft, 2021).

While some spike protein mutations have been associated with increased transmissibility or severity, the direct impact of many prevalent mutations on clinical manifestations remains unclear. Moreover, factors such as age and gender have been reported to affect COVID-19 outcomes, suggesting a complex interplay between viral genetics and host demographics. Understanding these relationships is crucial for developing effective public health strategies and personalized approaches to patient care.

Previous studies have utilized reverse vaccinology and computational methods to design multiepitope vaccines targeting major SARS-CoV-2 variants, aiming to enhance vaccine efficacy against emerging strains. Additionally, research into antiviral pharmacological strategies has identified key viral targets—including the main protease (Mpro), spike glycoprotein, and TMPRSS2 inhibitors—that are essential for therapeutic interventions. These efforts underscore the importance of continuous genomic surveillance and the need to correlate genetic variations with clinical outcomes.

Bangladesh, a densely populated country with significant international travel, has been particularly susceptible to the spread of COVID-19 (Chowdhury et al., 2020). The Sylhet region has reported a significant number of cases, yet there is limited data on the genetic diversity of SARS-CoV-2 and its clinical implications in this area. Given the global interconnectedness of the pandemic and the potential for regional variations to influence disease outcomes, a comprehensive analysis specific to this region is essential. (Chakraborty et al., 2020).

In this study, we aim to investigate the correlation between prevalent SARS-CoV-2 spike protein mutations—D614G, A570D, D1118H, A222V, and P681R—and clinical outcomes in patients from the Sylhet region of Bangladesh. We also examine how patient demographics, specifically age and gender, may influence these relationships. Our analysis seeks to elucidate whether these mutations have a direct impact on clinical manifestations and recovery rates or if host factors play a more significant role.

By providing insights into the genetic diversity of SARS-CoV-2 and its association with clinical outcomes and patient demographics, we aim to shed light on the genetic diversity and evolutionary dynamics of SARS-CoV-2 in the area, providing important insights for public health strategies (Akter et al. 2023; Shahab et al. 2023).

## Materials and Methods

### Whole-genome sequencing

SARS-CoV-2 detection via RT-PCR assay (Novel Coronavirus (2019-nCoV) Nucleic Acid Diagnostic Kit, Sansure Biotech) from nasopharyngeal specimens of 37 COVID- 19-positive patients collected from Sylhet, Bangladesh. Library preparation and next generation sequencing of the samples were conducted by the Genomic Research Laboratory of BCSIR. All methods were performed in accordance with the relevant guidelines and regulations. Informed consent was obtained from all subjects and/ or their legal guardian(s). Viral RNA was extracted using PureLink™ Viral RNA/DNA Mini Kit (Cat. no: 12280050, Thermofisher Scientific, USA)1. The cDNA of all samples was used to prepare paired-end libraries with the Nextera ™ DNA FlexLibrary Preparation kit according to the manufacturer’s instructions (Illumina Inc., San Diego, CA). The Library pool of the samples was sequenced using the S4 flow cell of Illumina NextSeq 550 instrument in a paired-end fashion (read length 151 bp; Illumina Inc.). FASTQ files were generated in the Illumina BaseSpace Sequence Hub (https://basespace.illumina.com). The generated data for after NGS sequencing were submitted to the GISAID database. The generated viral sequences were submitted to the GISAID database under accession numbers EPI_ISL_1538415, EPI_ISL_1538416, EPI_ISL_1538417, and so on Genomes of SARS-CoV-2 viruses were assembled using Basespace DRAGEN RNA Pathogen Detection V3.5.14 software with default parameters. Consensus FASTA files of the SARS-CoV-2 genome were generated using Basespace Dragon RNA Pathogen Detection software version 3.5.1 (https://basespace.illumina.com) with default settings. The sequences of SARS-CoV-2 isolates from Bangladeshi patients were compared to the reference SARS-CoV-2 sequence (NC_045512.1), through nucleotide substitutions. The positions of the nucleotides and amino acids were further confirmed from GeneBank reference sequences (NC_045512.1)

### Sequence Alignment and Editing

Sequences were aligned using the MAFFT algorithm, a widely recognized tool for multiple sequence alignment, which ensures high accuracy and efficiency (Katoh et al., 2019). Manual editing was conducted using the AliView program to verify alignment frames and correct potential errors. Nucleotide and amino acid positions were annotated starting from the 5’ UTR, with positions confirmed against the GeneBank reference sequence (NC_045512.1).

### Identification of Nucleotide Substitutions

Non-conserved nucleotide positions were identified, and the substitutions were analyzed for their potential effects on protein function using the AliView and MEGA- X software packages (Kumar et al., 2018). For this purpose, the nucleotide sequences were translated into amino acids, and the substitutions were assigned to specific protein-coding regions.

### Patients and Data Sources Cases Series of COVID-19

The case series analysis covers 37 patients with COVID-19 who were treated on site by the hospital’s medical team from January 29, 2020 to February 15, 2020.

### Data Set of COVID-19

The National Health Committee of the People’s Republic of Bangladesh introduced the diagnostic and clinical classification criteria and treatment plan (trial version 5) of COVID-19. The clinical classification of severity is as follows: (1) *Mild*, only mild symptoms, imaging shows no pneumonia; (2) *Moderate*, with fever, respiratory tract symptoms, and imaging shows pneumonia; (3) *Severe*, if any of the following signs are present: (a) Respiratory distress, respiratory rate ≥ 30 beats / min; (b) At rest, finger oxygen saturation ≤ 93% (c) arterial blood oxygen partial pressure (PaO_2_/oxygen concentration (FiO_2_) ≤ 300 mmHg (1 mmHg = 0.133 kPa); (4) *Critical*, one of the following conditions: (a) respiratory failure occurs and requires mechanical ventilation, (b) Shock occurs, (c) admission to ICU is required due to multiple organ failure.

### Statistical Analysis

We employed descriptive statistics to summarize the demographic and clinical data. Student’s t-test was used to compare differences between groups, with statistical significance set at P < 0.05. We performed all statistical analyses using R Studio software version 4.2.1, which offers robust tools for data analysis and visualization (Team, 2020).

## Results

### Case Series of Covid-19

To better understand the data, we have created a table to explain the description. The values of the individual characteristics are listed in **Table 1**.

**Table 1:** Overview of Patient Features, Clinical Symptoms, and SARS-CoV-2 Spike Mutations in Sylhet Region, Bangladesh.

| Feature | Type | Value |
| --- | --- | --- |
| age | Numerical | 5 - 75 |
| sex | Nominal | (0)female, (1)male |
| Location | Nominal | (1) Sylhet, (2) Habiganj, (3) Moulvibazer, (4) `Sunamganj |
| Days.since.last.sampling | Numerical | 29 - 60 |
| Lack.of.smell | Nominal | (0)No, (1)Yes |
| Diarrhoea | Nominal | 0)No, (1)Yes |
| Recovered | Nominal | 0)No, (1)Yes |
| Hospitalized | Nominal | 0)No, (1)Yes |
| Spike.D614G | Nominal | (0)Absent (1)Present |
| Spike.A570D | Nominal | (0)Absent (1)Present |
| Spike.D1118H | Nominal | (0)Absent (1)Present |
| Spike.A222V | Nominal | (0)Absent (1)Present |
| Spike.P681R | Nominal | (0)Absent (1)Present |
| Clade | Nominal | (1) GR, (2) GRY, (3) GH, (4) GV, (5) G |

The dataset contains information on patients with COVID-19 from different locations in the Sylhet region of Bangladesh. It contains both numeric and nominal characteristics. Age (between 5 and 75 years) and days since last sampling (29 to 60 days) are the numerical characteristics. Nominal characteristics include gender (0 for female, 1 for male), location (1 for Sylhet, 2 for Habiganj, 3 for Moulvibazar, 4 for Sunamganj) and clinical symptoms such as lack of smell (anosmia) and diarrhea (0 for no, 1 for yes). In addition, the dataset records whether patients have recovered, have been hospitalized, and whether certain spike protein mutations (D614G, A570D, D1118H, A222V, P681R) are present (0 for not present, 1 for present). Finally, the dataset classifies patients into one of five clades (1: GR, 2: GRY, 3: GH, 4: GV, 5: G) representing different SARS-CoV-2 lineages. This structured dataset allows for a comprehensive analysis of the correlation between patient characteristics, symptoms, and spike protein mutations in the context of specific clades and geographical locations.

### Observational Study Type

The data indicates that out of the total sample, 22 patients were not hospitalized due to COVID-19, while 15 patients required hospitalization. The dependent variable in this study is hospitalization status, which is qualitative in nature. The independent variables in this study include age, gender, and other relevant factors.

Regarding gender distribution, 37.8% of the patients were female, and 62.2% were male (Figure 1). The analysis of hospitalization based on gender revealed that, among females, 9 out of 14 (64%) were hospitalized due to COVID-19, while 13 out of 23 males (57%) were hospitalized. These results suggest a slightly higher proportion of hospitalization among females compared to males in this dataset.

**Figure 1:**
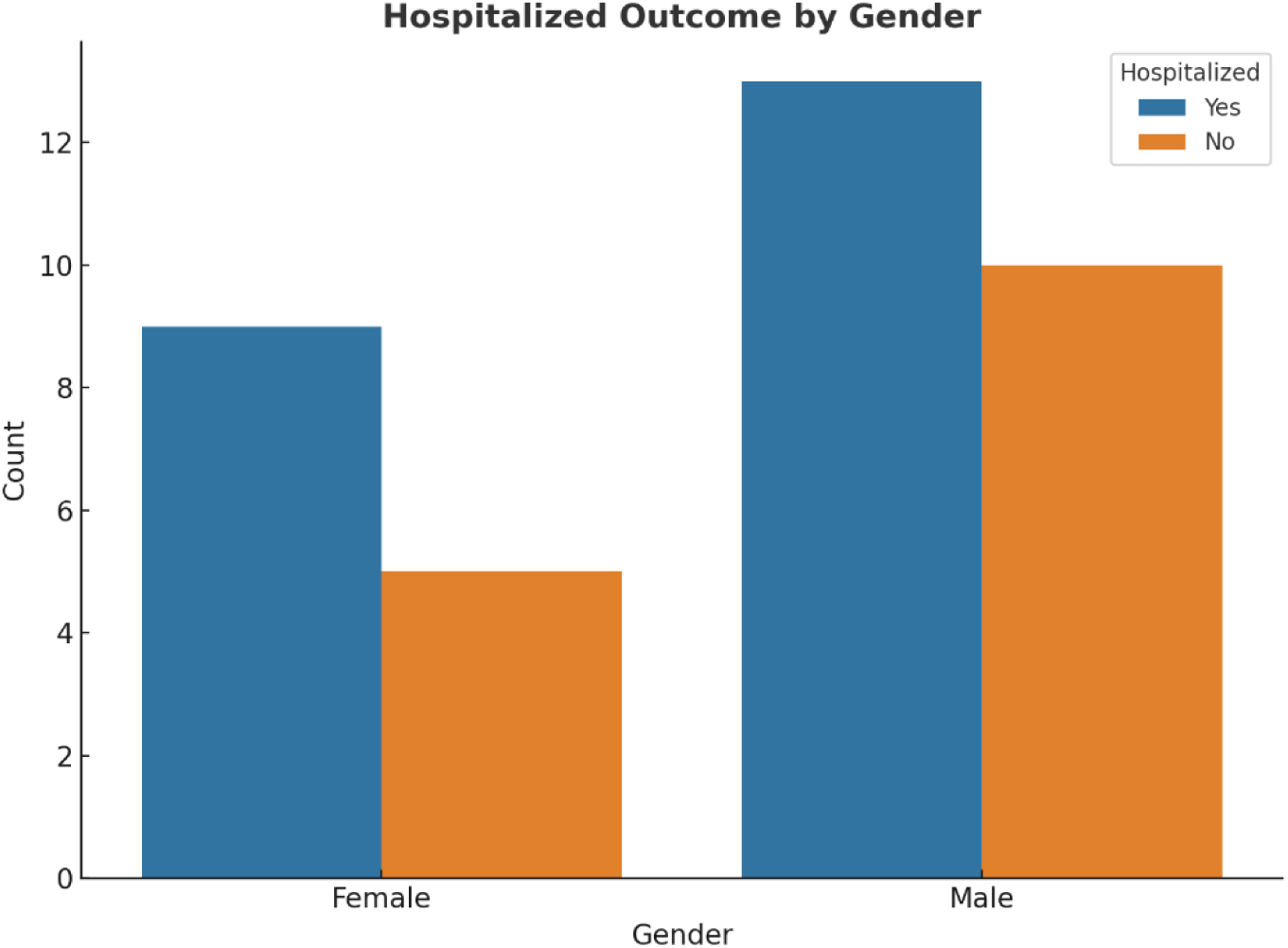
Gender-Based Comparison of COVID-19 Hospitalization Rates

**Figure 2:**
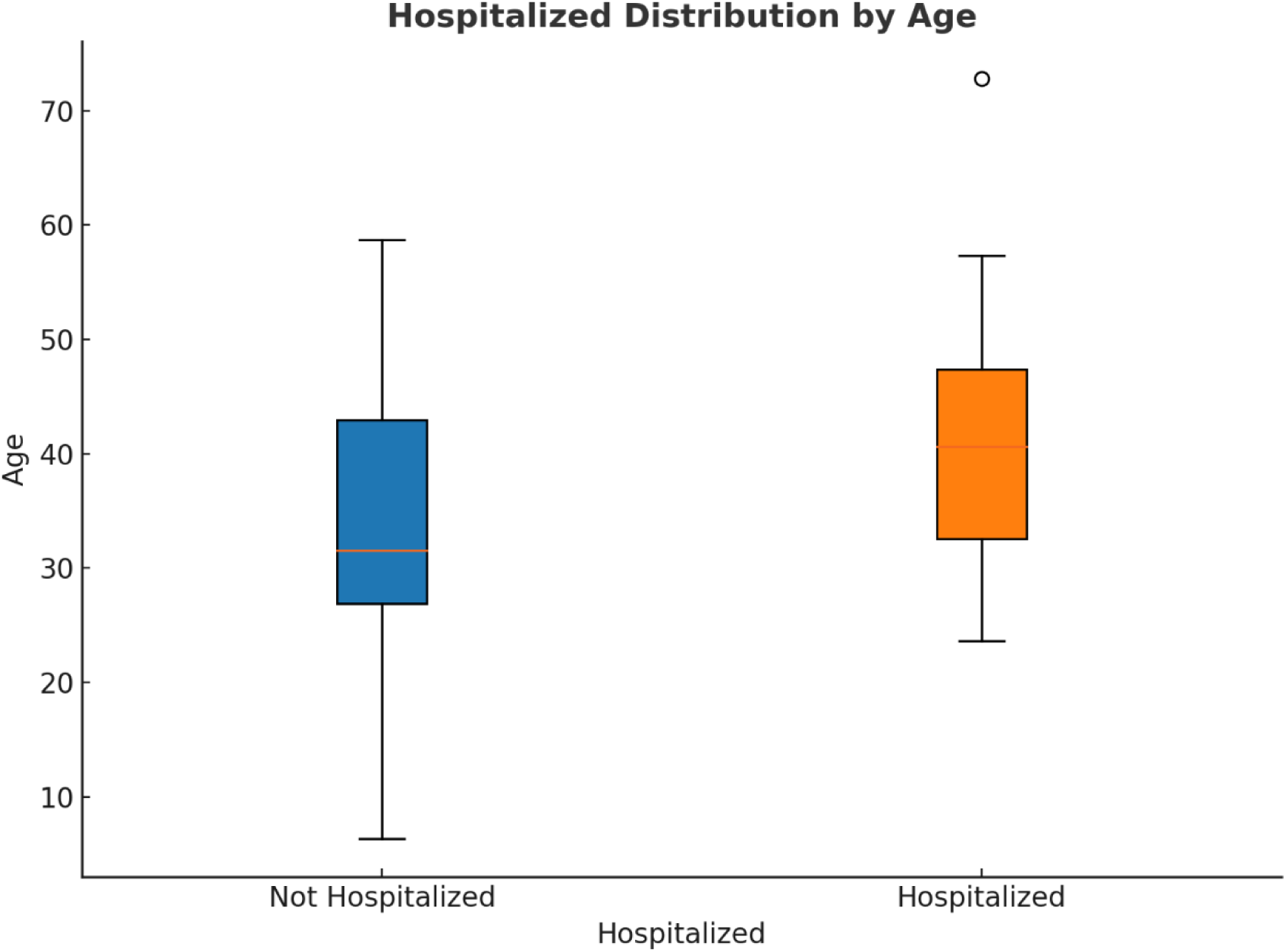
Age Distribution of Hospitalized vs. Non-Hospitalized COVID-19 Patients. The boxplot illustrates the age range within each group. Hospitalized patients tend to have a higher median age compared to non-hospitalized patients, as shown by the interquartile range. An outlier is observed in the hospitalized group.

Figure 3 illustrates the normal distribution of the age data analyzed, with the probability density and cumulative probability displayed. The probability density (blue curve) describes the relative frequency of different age groups, peaking near the population mean. The cumulative probability curve (orange curve) shows that approximately 73.8% of individuals are aged 50 years or younger, highlighting the concentration of younger individuals in the dataset.

**Figure 3:**
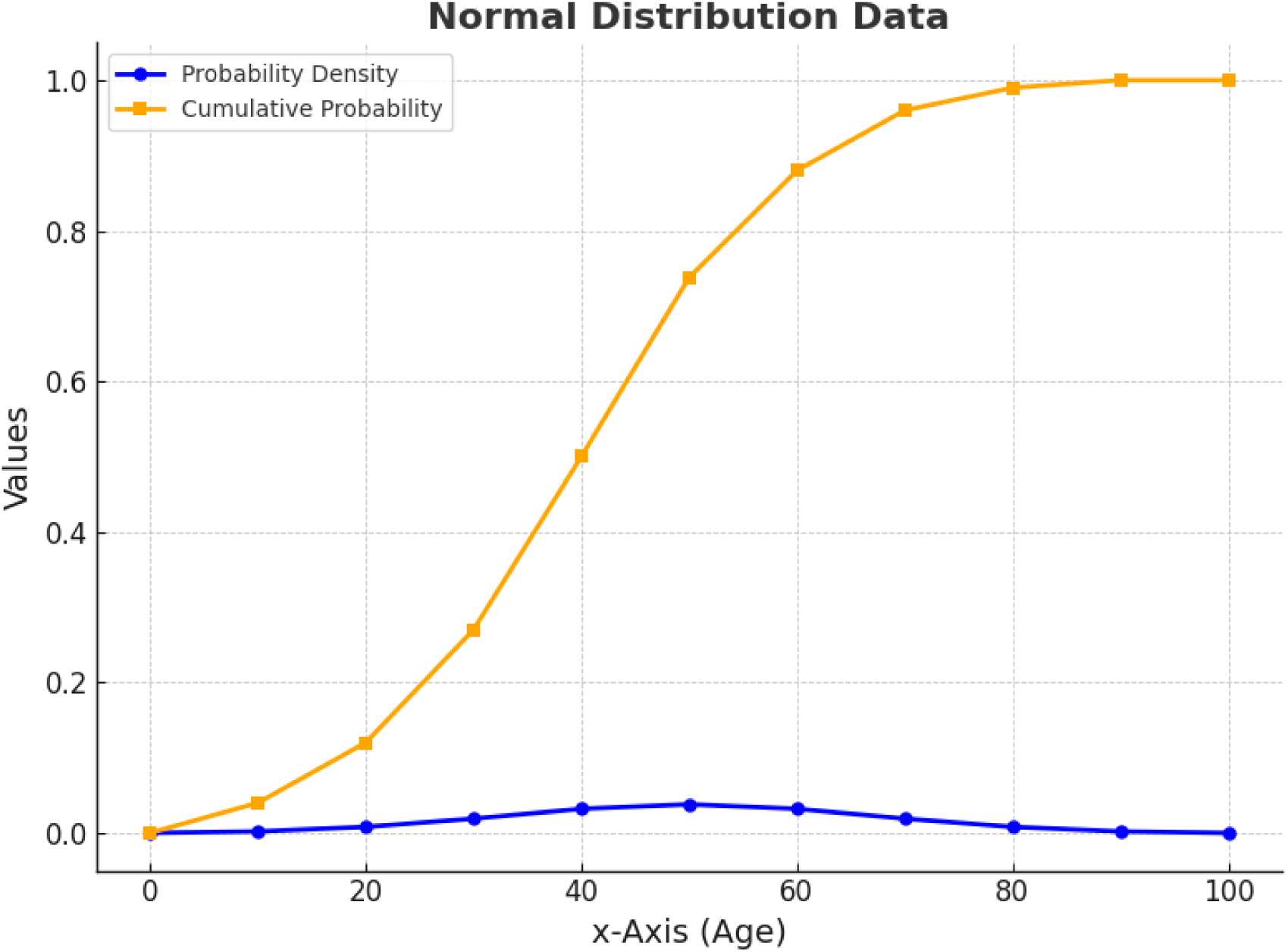
Graphical representation of the probability density and cumulative probability in relation to age. The blue curve represents the probability density for different age groups, while the orange curve indicates the cumulative probability, showing the proportion of individuals up to a certain age.

In Figure 4, the distribution is further broken down by gender, revealing distinct differences in age between male and female patients. Female patients tend to be younger, with a mean age of 31.20, while male patients exhibit a broader age range with a mean age of 42.17. This distinction suggests that male patients in the dataset are generally older, which may indicate a higher risk of severe outcomes or hospitalization in older male individuals compared to younger female patients. Together, these figures underscore the age-related patterns in the dataset and provide insights into how gender and age may influence COVID-19 severity.

**Figure 4:**
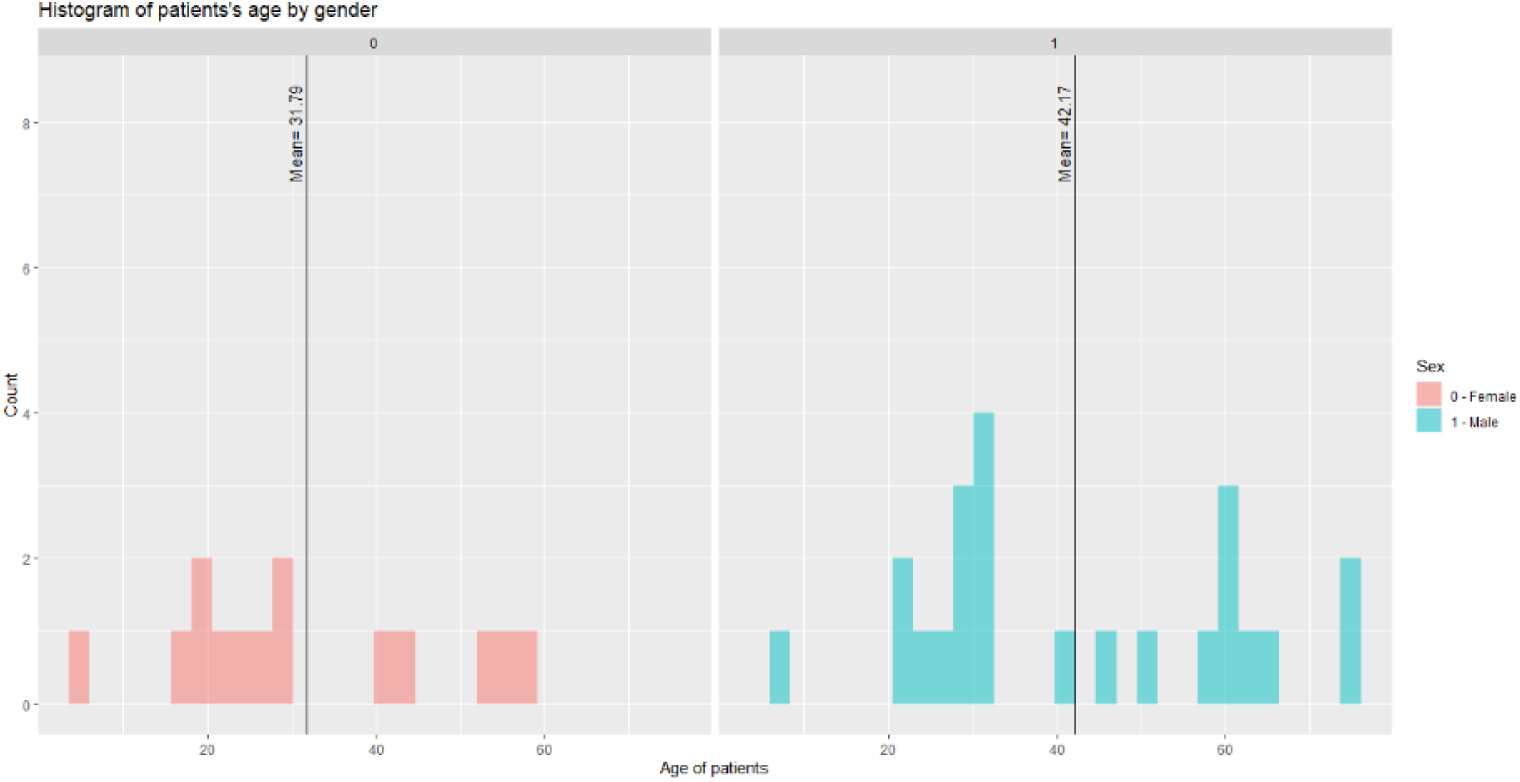
Age Distribution of Patients by Gender with Mean Age Indication

Figure 5 presents a comparative analysis of hospitalized COVID-19 patients based on two symptoms: diarrhoea (A) and lack of smell (B). In (A), the majority of hospitalized patients (90%) were affected by diarrhoea, while only 10% were not. In contrast, among non-hospitalized patients, a higher proportion (62.5%) experienced diarrhoea compared to 37.5% who did not. This suggests a potential association between diarrhoea and the severity of COVID-19, reflected in hospitalization rates.

**Figure 5:**
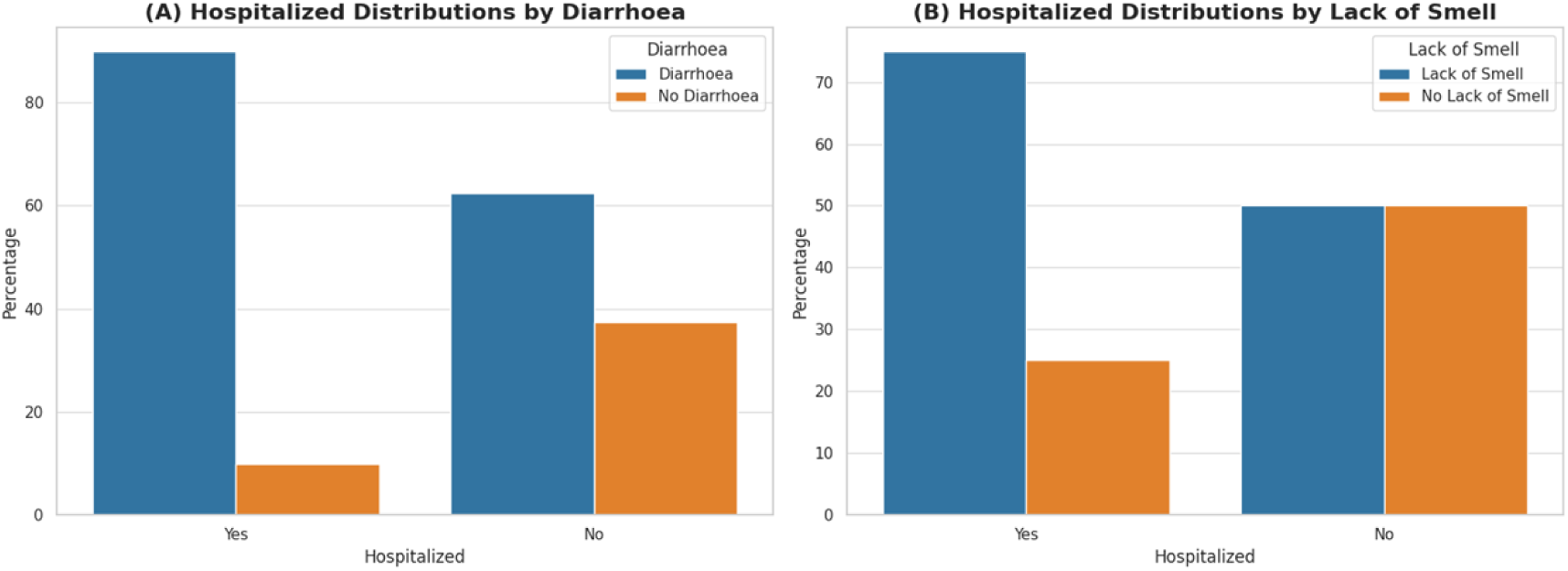
Comparison of Hospitalized COVID-19 Patients by Diarrhoea (A) and Lack of Smell (B)

In (B), the distribution of hospitalized patients based on the presence of smell loss shows that 75% of hospitalized patients experienced loss of smell, while 25% did not. Interestingly, the non-hospitalized group had an even split between those with and without smell loss (50% each). This suggests that loss of smell may not be as strongly correlated with the likelihood of hospitalization compared to diarrhoea. These findings highlight the varying impact of different symptoms on COVID-19 severity and hospital admission rates.

Figure 6 provides a correlation heatmap that visually represents the relationships between clinical variables, patient demographics, and spike protein mutations. The strength and direction of these correlations are indicated by the intensity of the colors— blue representing positive correlations and red representing negative correlations. Darker shades reflect stronger relationships.

**Figure 6:**
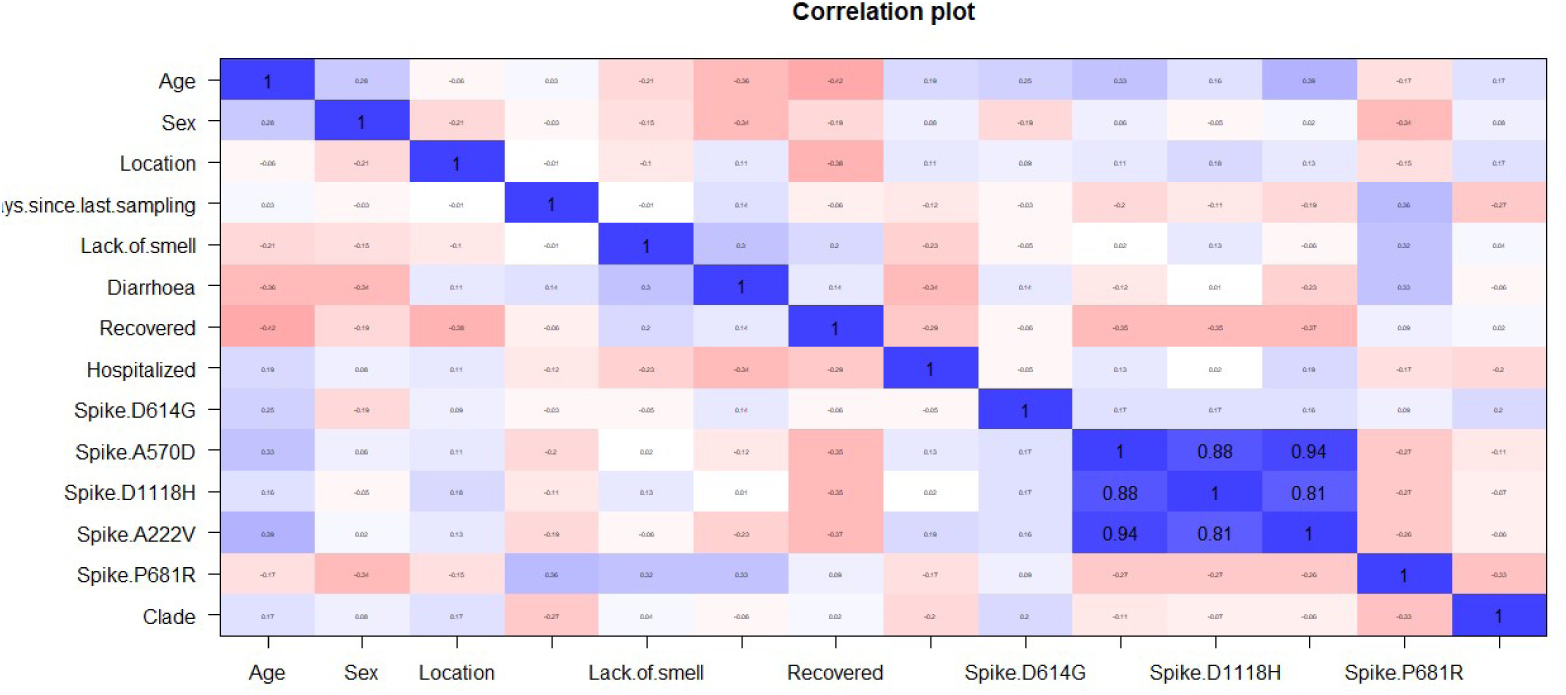
Correlation Heatmap Between Clinical Variables and Spike Mutations in COVID- 19 Patients

The most notable observation is the strong positive correlation between specific spike protein mutations. Spike A570D, D1118H, and A222V exhibit correlations above 0.9 (deep blue), suggesting that these mutations frequently co-occur within the dataset. This pattern may point to their association with certain SARS-CoV-2 variants, particularly variants of concern such as Alpha and Delta, which are known for increased transmissibility, immune evasion, or viral adaptation. The frequent co-occurrence of these mutations could indicate the presence of clusters of infections related to these variants.

Additionally, age demonstrates a moderate negative correlation with **recovery** (-0.42), indicating that older patients are less likely to recover within the same timeframe as younger patients. This finding aligns with established literature, which emphasizes the increased vulnerability of older individuals to severe outcomes and prolonged recovery times. Factors such as diminished immune responses and the presence of comorbidities likely contribute to these age-related differences in recovery rates.

A particularly interesting finding is the lack of strong correlations between spike mutations and clinical symptoms such as loss of smell or diarrhoea. Except for a moderate correlation with the P681R mutation (greater than 30%), these intersections exhibit correlations near zero. This suggests that the presence of these mutations does not strongly influence whether a patient experiences these symptoms. While spike mutations may affect the virus’s transmissibility or immune evasion, they do not appear to directly impact the presentation of these specific clinical symptoms in this patient population.

Figure 7 presents the feature importance for predicting outcomes among male and female COVID-19 patients using IncNodePurity, which measures the reduction in node impurity (i.e., the quality of a split in decision trees) across various features. This analysis highlights the relative importance of different clinical and genetic features in determining patient outcomes for each gender.

**Figure 7:**
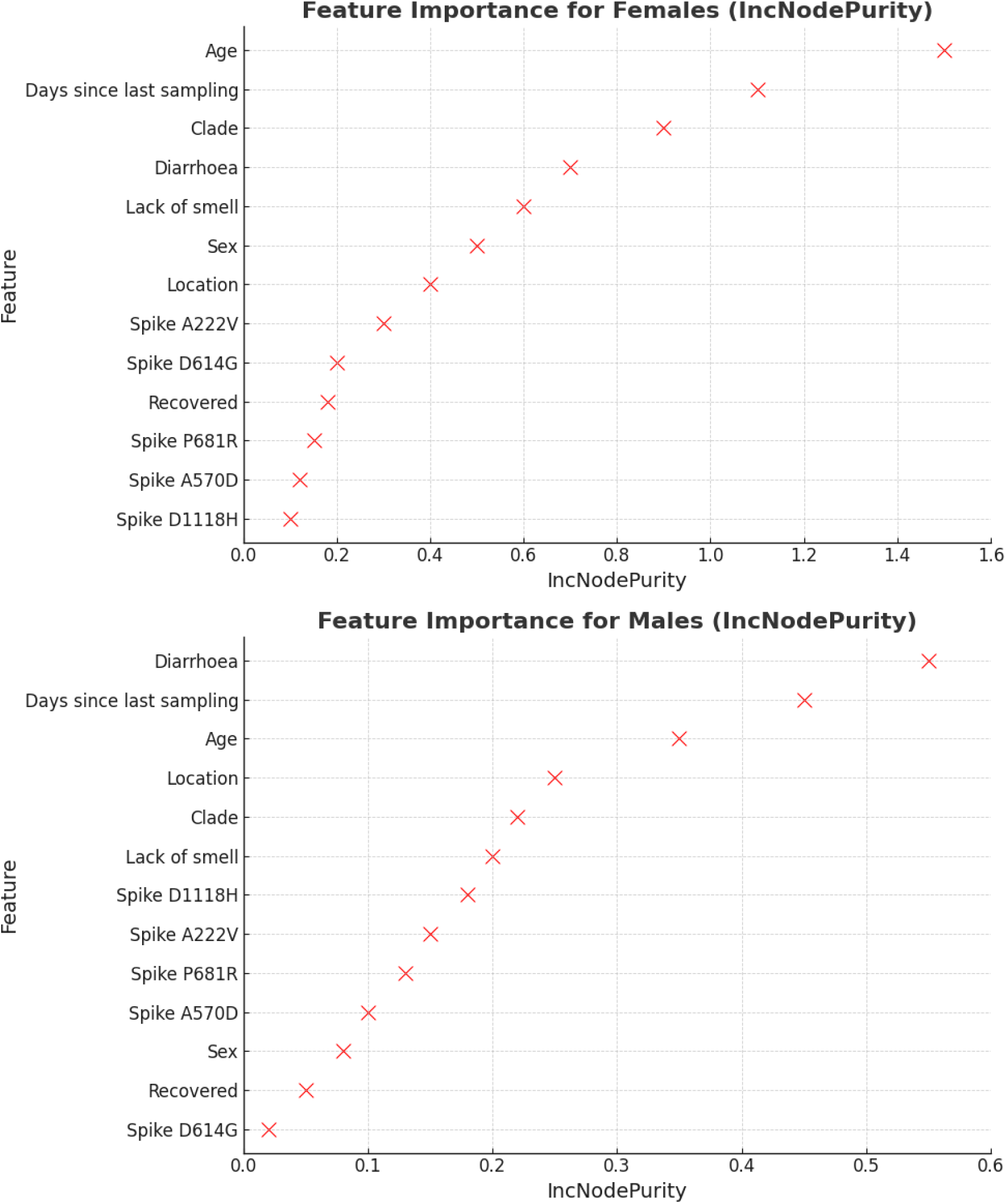
Gender-Specific Feature Importance for Predicting COVID-19 Outcomes Using IncNodePurit

The most influential feature for predicting outcomes in *male* patients is Diarrhoea, followed by Days Since Last Sampling and Age. This suggests that, among males, the presence of diarrhoea may be a significant indicator of disease severity or hospitalization. The number of days since the last sampling also plays a critical role, which may reflect disease progression or the timing of symptom onset. Furthermore, spike protein mutations, such as Spike D1118H and Spike A222V, also have some predictive power but are ranked lower compared to clinical features like age and location.

In contrast, *Age* is the most significant predictor of outcomes in female patients, showing much higher importance than in males. This suggests that age is a stronger factor in determining disease outcomes for females, likely due to age-related factors such as immune system function and comorbidities. Days Since Last Sampling and Clade follow closely, indicating that these features also have an important role in predicting the progression of the disease in female patients. Clinical symptoms like Diarrhoea and Loss of Smell are less prominent but still contribute to the overall prediction.

A notable difference between the two groups is the relative importance of spike protein mutations. For males, mutations like Spike D1118H and Spike P681R rank higher in feature importance, suggesting that genetic variations may have a larger impact on male patients. In contrast, these mutations are less significant for female patients, where Age and clinical features take precedence.

This gender difference in feature importance could reflect underlying biological differences in how male and female patients respond to SARS-CoV-2, or it may indicate that different factors influence disease severity across genders.

Table 2 shows that the spike D614G mutation was found in 94.6% of the patients in the data set. In terms of age, 43.2% of patients aged 0-30 years had the mutation, while 18.9% were between 31-50 years old and 32.4% were 51 years and older. In addition, all patients who reported diarrhea (24.3% of the total population) tested positive for spike D614G, while of the patients with loss of smell (40.5% of the total population), 37.8% carried the mutation. Despite these distributions, no statistically significant association was found between the Spike D614G mutation and gender, age, diarrhea, or odor loss, as determined by Fisher’s exact test.

**Table 2:** A Cross Tabulation of Spike D614G and some selected variables.

| Variables |  | Spike D614G (N, %) |  | Total | p-value |
| --- | --- | --- | --- | --- | --- |
|  |  | No (0) | Yes (1) |  |  |
| Total |  | 2 (5.4) | 35 (94.6) | 37<br>(100.0) |  |
| Sex | Female | 0 (0.0) | 14 (37.8) | 14 (37.8) | 0.517 |
|  | Male | 2 (5.4) | 21 (56.8) | 23 (62.2) |  |
| Age | 0 - 30 | 1 (2.7) | 16 (43.2) | 17 (45.9) | 0.657 |
|  | 31 - 50 | 1 (2.7) | 7 (18.9) | 8 (21.6) |  |
|  | 51 and above | 0 (0.0) | 12 (32.4) | 12 (32.4) |  |
| Diarrhea | No | 2 (5.4) | 26 (70.3) | 28(75.6) | 1.000 |
|  | Yes | 0 (0.0) | 9 (24.3) | 9 (24.3) |  |
| Lack of Smell | No | 1 (2.7) | 21 (56.8) | 22 (59.5) | 1.000 |
|  | Yes | 1 (2.7) | 14 (37.8) | 15 (40.5) |  |
| Hospitalized | No (R) | 1 (5.6) | 17 (94.4) | 18 (58.1) | 0.6474 |
|  | Yes | 1 (7.7) | 12 (92.3) | 13 (41.9) |  |

Similarly, Table 3 reports the prevalence of the spike A570D mutation, which was found in 32.4% of patients (10.8% were female and 21.6% were male). The distribution of the mutation across age groups showed that 13.5% of patients in the age group 0-30 years had the mutation, 2.7% in the group 31-50 years and 16.2% in the group 51 years and older. In terms of clinical symptoms, 5.4% and 13.5% of patients tested positive for spike A570D and had diarrhea and loss of smell, respectively. As with Spike D614G, no statistically significant association was found between the Spike A570D mutation and the selected variables of gender, age, diarrhea or loss of smell (Fisher’s exact test).

**Table 3:** A Cross Tabulation of Spike A570D and some selected variables.

| Variables |  | Spike A570D (N, %) |  | Total | p-value |
| --- | --- | --- | --- | --- | --- |
|  |  | No (0) | Yes (1) |  |  |
| Total |  | 25 (67.5) | 12 (32.4) | 37<br>(100.0) |  |
| Sex | Female | 10 (27.0) | 4 (10.8) | 14 (37.8) | 1.000 |
|  | Male | 15 (40.5) | 8 (21.6) | 23 (62.2) |  |
| Age | 0 - 30 | 12 (32.4) | 5 (13.5) | 17 (45.9) | 0.633 |
|  | 31 - 50 | 7 (18.9) | 1 (2.7) | 8 (21.6) |  |
|  | 51 and above | 6 (16.2) | 6 (16.2) | 12 (32.4) |  |
| Diarrhea | No | 18 (48.6) | 10 (27.0) | 28(75.6) | 0.687 |
|  | Yes | 7 (18.9) | 2 (5.4) | 9 (24.3) |  |
|  |  | No (0) | Yes (1) |  |  |
| Lack of Smell | No | 15 (40.5) | 7 (18.9) | 22 (59.5) | 1.000 |
|  | Yes | 10 (27.0) | 5 (13.5) | 15 (40.5) |  |
| Hospitalized | No | 12 (66.7) | 6 (33.3) | 18 (58.1) | 0.7324 |
|  | Yes | 8 (61.5) | 5 (38.5) | 13 (41.9) |  |

According to Table 4, the spike D1118H mutation was found in 32.4% of the patients in the dataset (13.5% were female and 18.9% were male). In terms of age distribution, 16.2%, 2.7% and 13.5% of patients with mutation were in the 0-30, 31-50 and 51+ age groups, respectively. In terms of clinical symptoms, only 8.1% of patients had diarrhea and tested positive for spike D1118H, and 16.2% of patients reported loss of smell and carried the mutation. Again, no statistically significant association was found between spike D1118H and gender, age, diarrhea or loss of smell based on Fisher’s exact test.

**Table 4:** A Cross Tabulation of Spike D1118H and some selected variables.

| Variables |  | Spike D1118H (N, %) |  | Total | p-value |
| --- | --- | --- | --- | --- | --- |
|  |  | No (0) | Yes (1) |  |  |
| Total |  | 25 (67.5) | 12 (32.4) | 37 (100.0) |  |
| Sex | Female | 9 (24.3) | 5 (13.5) | 14 (37.8) | 1.000 |
|  | Male | 16 (43.2) | 7 (18.9) | 23 (62.2) |  |
| Age | 0 - 30 | 11 (29.7) | 6 (16.2) | 17 (45.9) | 0.359 |
|  | 31 - 50 | 7 (18.9) | 1 (2.7) | 8 (21.6) |  |
|  | 51 and above | 7 (18.9) | 5 (13.5) | 12 (32.4) |  |
| Diarrhea | No | 19 (51.4) | 9 (24.3) | 28 (75.6) | 1.000 |
|  | Yes | 6 (16.2) | 3 (8.1) | 9 (24.3) |  |
| Lack of Smell | No | 16 (43.2) | 6 (16.2) | 22 (59.5) | 0.488 |
|  | Yes | 9 (24.3) | 6 (16.2) | 15 (40.5) |  |
| Hospitalized | No | 11 (61.1) | 7 (38.9) | 18 (58.1) | 0.7213 |
|  | Yes | 9 (69.2) | 4 (30.8) | 13 (41.9) |  |

Table 5 shows that the spike A222V mutation was present in 29.7% of patients. Among these patients, 10.8% were female and 18.9% were male. In terms of age distribution, 10.8% of patients were 0-30 years old and tested positive for spike A222V, 2.7% were in the 31-50 years group, and 16.2% were in the 51 years and older group. Of the patients, 2.7% reported diarrhea and carried the mutation, while 18.9% of patients had loss of smell and tested positive for spike A222V. No statistically significant association was found between spike A222V and any of the selected variables, as shown by Fisher’s exact test.

**Table 5:** A Cross Tabulation of Spike A222V and some selected variables.

| Variables |  | Spike A222V (N, %) |  | Total | p-value |
| --- | --- | --- | --- | --- | --- |
|  |  | No (0) | Yes (1) |  |  |
| Total |  | 26 (70.3) | 11 (29.7) | 37<br>(100.0) |  |
| Sex | Female | 10 (27.0) | 4 (10.8) | 14 (37.8) | 1.000 |
|  | Male | 16 (43.2) | 7 (18.9) | 23 (62.2) |  |
| Age | 0 - 30 | 13 (35.1) | 4 (10.8) | 17 (45.9) | 0.883 |
|  | 31 - 50 | 7 (18.9) | 1 (2.7) | 8 (21.6) |  |
|  | 51 and above | 6 (16.2) | 6 (16.2) | 12 (32.4) |  |
| Diarrhea | No | 18 (48.6) | 10 (27.0) | 28(75.6) | 0.229 |
|  | Yes | 8 (21.6) | 1 (2.7) | 9 (24.3) |  |
| Lack of Smell | No | 15 (40.5) | 7 (18.9) | 22 (59.5) | 1.000 |
|  | Yes | 11 (29.7) | 4 (10.8) | 15 (40.5) |  |
| Hospitalized | No | 13 (72.2) | 5 (27.8) | 18 (58.1) | 0.7040 |
|  | Yes | 8 (61.5) | 5 (38.5) | 13 (41.9) |  |

Table 6 shows that spike P681R was present in 13.5% of patients. Spike P681R was rarely found in the patients studied (10.8% were women and 2.7% were men). Again, 4 of the 5 individuals who tested positive for spike P681R had a lack of odor. Similarly, 3 of the 5 individuals who tested positive for spike P681R had diarrhea. In terms of assessing the statistical association, there was no association between spike P681R and the selected variables (gender, age, diarrhea, and lack of odor) from the Fishers exact test.

**Table 6:**
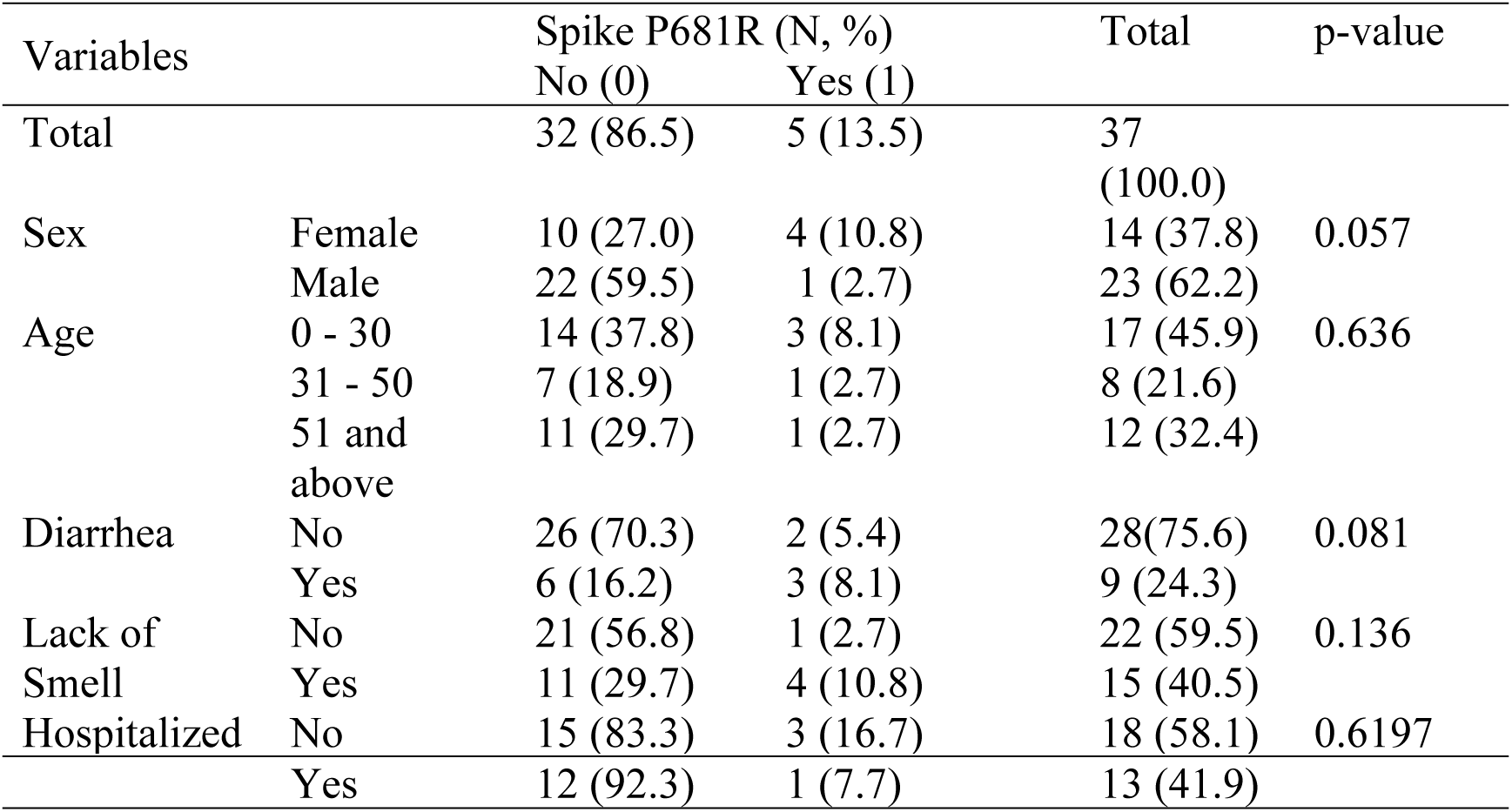
A Cross Tabulation of Spike P681R and some selected variables.

Table 7 presents the frequency and percentage distribution of key demographic and clinical variables in the dataset. Regarding clinical symptoms, 59.5% of the patients did not report a loss of smell, while 40.5% did. In terms of gastrointestinal symptoms, 75.7% of patients did not experience diarrhea, whereas 24.3% did. The data on hospitalization status shows that out of the 31 patients for whom hospitalization data was available, 58.1% were not hospitalized, while 41.9% required hospitalization. This provides an overview of the distribution of symptoms and hospitalization status in the study population, highlighting the higher prevalence of male patients and the relatively low incidence of diarrhea compared to loss of smell.

**Table 7:** Frequency Distribution Table.

| Variables |  | Frequency | Percentage (%) |
| --- | --- | --- | --- |
| Sex | Female | 14 | 37.8 |
|  | Male | 23 | 62.2 |
|  | Total | 37 | 100.0 |
| Lack of smell | No | 22 | 59.5 |
|  | Yes | 15 | 40.5 |
|  | Total | 37 | 100.0 |
| Diarrhea | No | 28 | 75.7 |
|  | Yes | 9 | 24.3 |
|  | Total | 37 | 100.0 |
| Hospitalized | No | 18 | 58.1 |
|  | Yes | 13 | 41.9 |
|  | Total | 31 | 100.0 |

The bar charts display the prevalence of various spike protein mutations among the patients examined (Figure 8). The D614G mutation was present in the majority of the population, with 94.59% (n = 35) testing positive and only 5.41% (n = 2) negative, indicating its dominance in this sample. For the A570D mutation, most patients (67.57%, n = 25) tested negative, while 32.43% (n = 12) carried the mutation. Similarly, the D1118H mutation was absent in 67.57% (n = 25) of patients, with 32.43% (n = 12) testing positive for this variant. The A222V mutation showed a similar pattern, with 70.27% (n = 26) of patients testing negative and 29.73% (n = 11) positive. Finally, the P681R mutation was the least common, with 86.49% (n = 32) of patients testing negative and 13.51% (n = 5) carrying the mutation.

**Figure 8:**
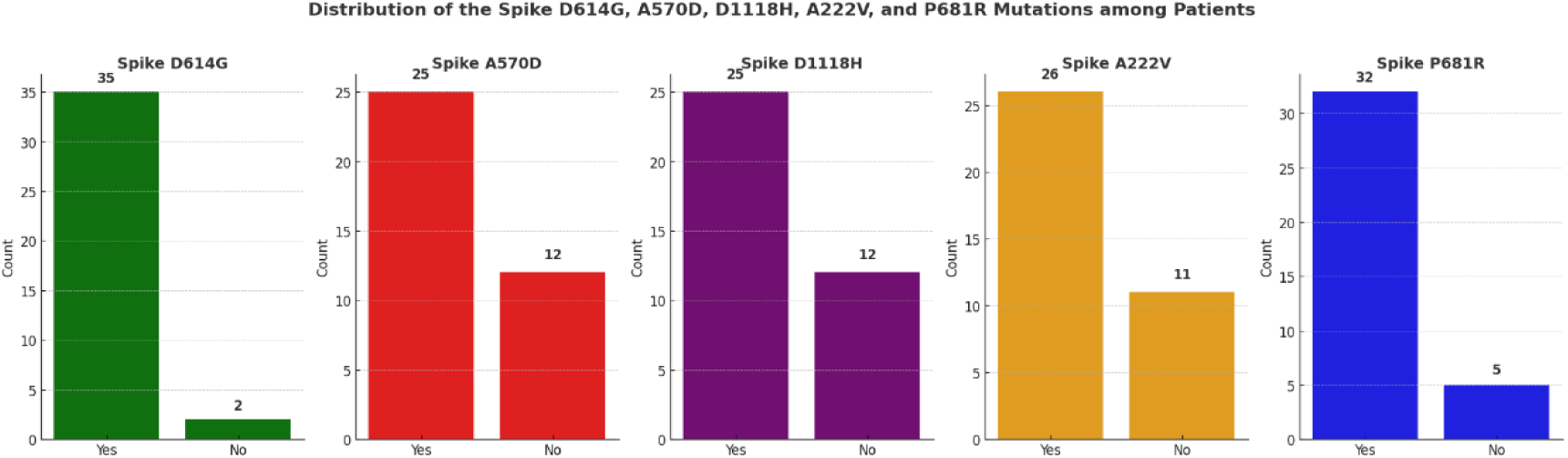
Distribution of the Spike D614G, A570D, D1118H, A222V and P681R Mutation among Patients

**Fig 9:**
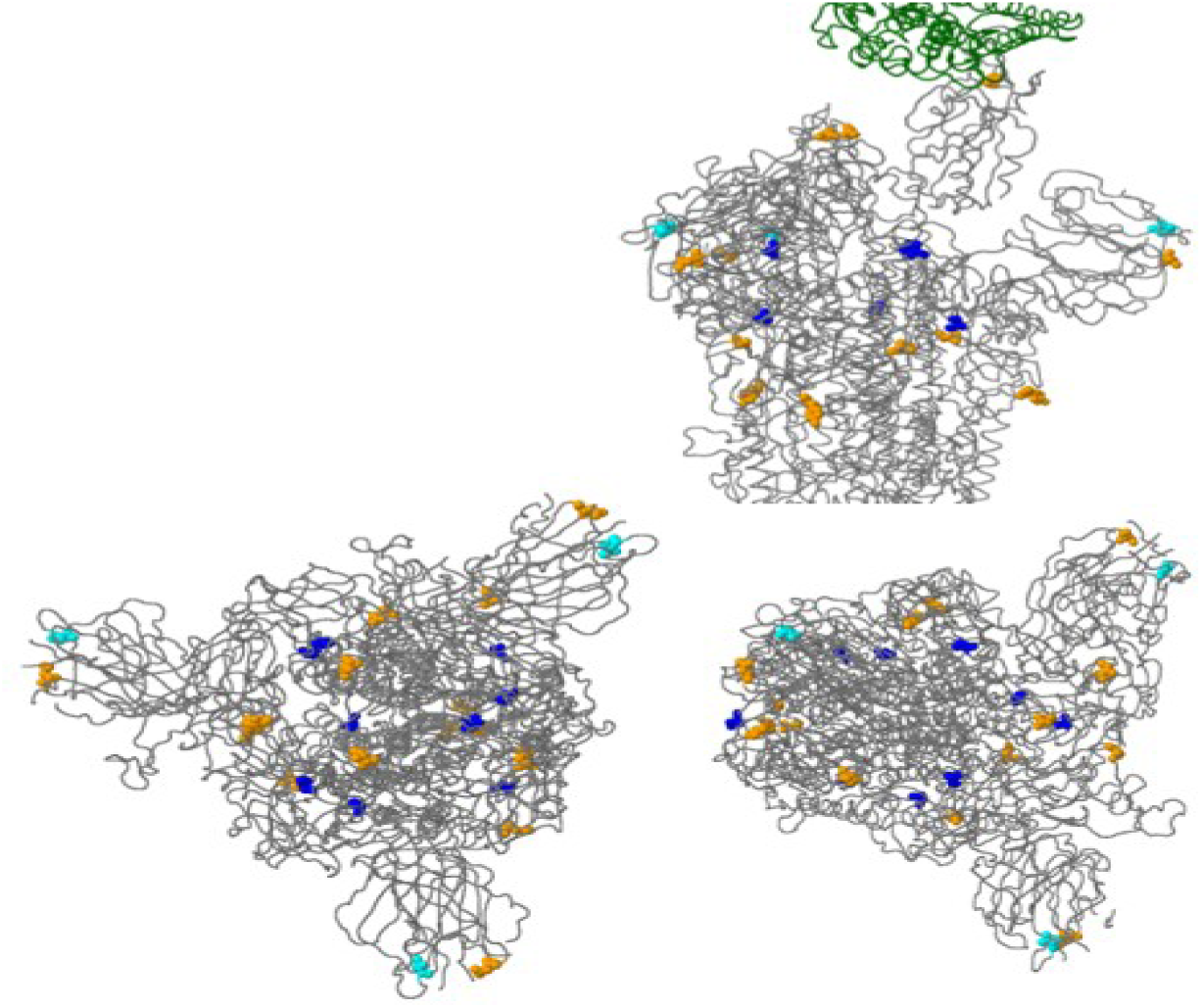
Distribution pattern of mutation within the spike proteins of SARS-CoV-2

The Fisher’s exact test results show that most spike variants (D614G, A570D, D1118H, A222V) do not exhibit significant associations with the selected variables (sex, lack of smell, and diarrhea), indicated by high p-values. However, the P681R variant shows a marginal association with sex (p = 0.0574) and slightly lower p-values for lack of smell (p = 0.1362) and diarrhea (p = 0.0812), suggesting potential trends that might require further investigation with a larger sample size (Table 8).

**Table 8:** Test of Association between the various spikes and some selected variables using Fishers’ exact test.

|  | Sex | Lack of smell | Diarrhea |
| --- | --- | --- | --- |
| Spike D614G | 0.5165 | 1.000 | 1.000 |
| Spike A570D | 1.000 | 1.000 | 0.687 |
| Spike D1118H | 1.000 | 0.4879 | 1.000 |
| Spike A222V | 1.000 | 1.000 | 0.2293 |
| Spike P681R | 0.0574 | 0.1362 | 0.0812 |

### Identification of Nucleotide Substitutions

We identified several common nucleotide substitutions in the Sylhet sequences, with C3037T, C14408T, and A23403G being the most prevalent. These mutations are part of a haplotype frequently observed in European strains, suggesting a possible introduction from Europe. Other notable substitutions included C241T, C1059T, and G25563T. The full list of identified substitutions and their corresponding protein-coding regions is presented in Table 9.

**Table 9:** Nucleotide Substitutions and Their Effects on Protein-Coding Regions in SARS-CoV-2 Genome.

| Nucleotide Position | Reference Nucleotide | Mutant Nucleotide | Protein-Coding Region | Effect on Amino Acid |
| --- | --- | --- | --- | --- |
| C14408T | C | T | ORF1b | P4715L |
| A23403G | A | G | Spike | D614G |
| C241T | C | T | 5' UTR | one |
| C1059T | C | T | ORF1a | T85I |
| G25563T | G | T | ORF3a | Q57H |

### Structural Analysis of Mutations in the SARS-CoV-2 Spike Protein

The structural analysis of the SARS-CoV-2 spike protein was conducted to elucidate the spatial distribution and potential impact of specific mutations. Figures 1, 2, and 3 depict the three-dimensional structure of the spike protein with key mutations highlighted (Fig 16). Figure 16 illustrates the three-dimensional structure of the spike protein, with notable mutations represented by color-coded spheres. The orange spheres denote specific amino acid residues of interest, potentially representing mutations that alter the protein’s function or interaction with host cells. The blue spheres indicate additional significant mutations or conserved regions that may play crucial roles in the protein’s stability or immune recognition. Cyan spheres represent other critical residues, possibly involved in binding or structural integrity. This figure provides a comprehensive view of how these mutations are spatially distributed within the protein.

These figures collectively provide a detailed structural analysis of the SARS-CoV-2 spike protein, highlighting the positions and potential impacts of various mutations. The spatial distribution of these mutations, as indicated by the color-coded spheres, suggests possible changes in the protein’s function, stability, and interaction with host cells or the immune system. The inclusion of interacting proteins in the structural analysis emphasizes the importance of these mutations in viral pathogenesis and potential therapeutic interventions. Understanding the structural implications of these mutations is crucial for guiding vaccine design and developing effective therapeutic strategies against SARS-CoV-2.

### Global Phylogenetic Analysis and Spike Protein Mutation Patterns in SARS-CoV- 2 Sequences

The figure 10 demonstrates the phylogenetic relationships and mutation distribution patterns of SARS-CoV-2 sequences, using maximum likelihood analysis. Panel (a) displays a phylogenetic tree with branch lengths representing genetic distance (mutations). The red dot marks the uploaded sequence, showing its relationship to other global sequences. Panel (b) illustrates the collection dates of genomes within the various tip groups, color-coded by country, with the United Kingdom dominating the dataset. Panel (c) focuses on the frequency of different spike protein mutations across the sequences, indicating that certain mutations, like D614G, are more frequent. Finally, panel (d) highlights lineage distribution (B.1.1.7 and B.1) based on collection dates, providing insights into how these variants have evolved over time. This comprehensive visual aids in understanding the genetic divergence, global distribution, and mutation patterns of SARS-CoV-2 variants.

**Fig 10:**
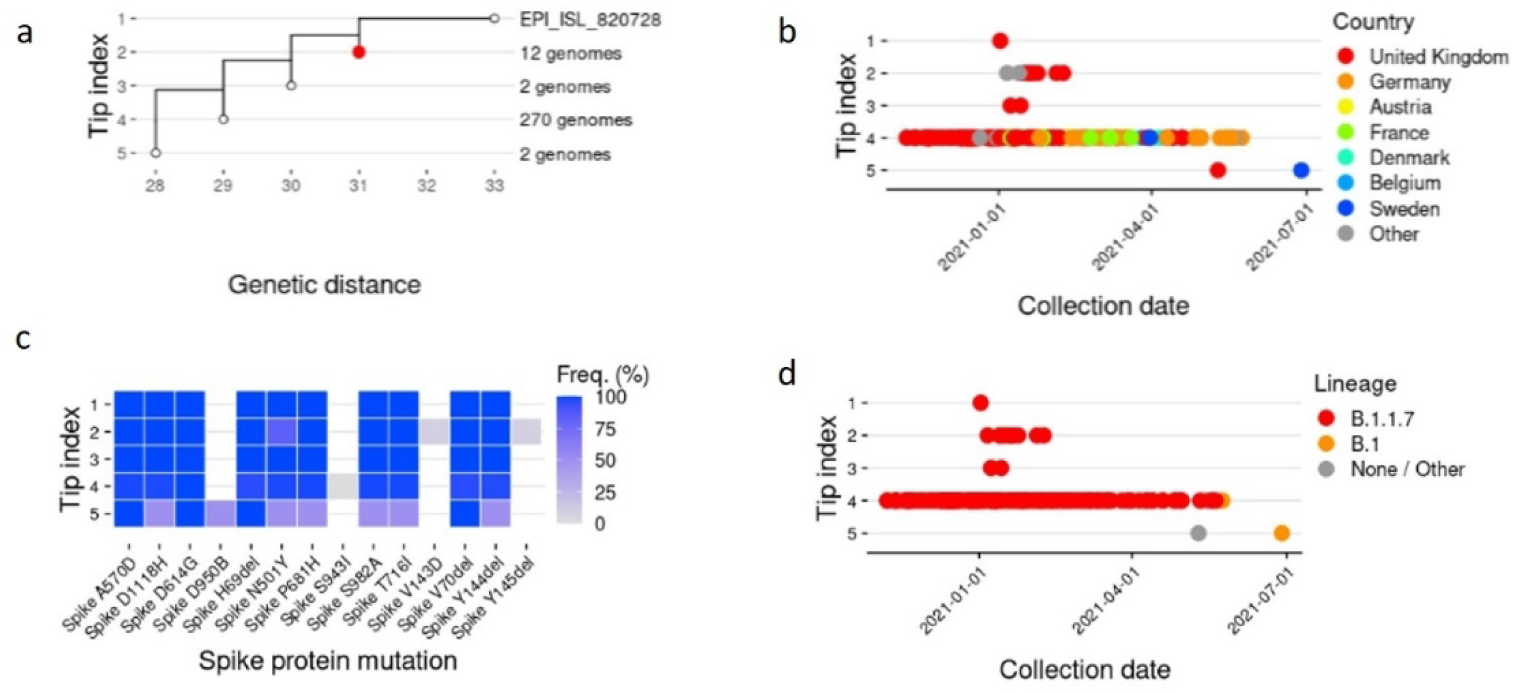
A maximum likelihood tree of the uploaded sequence, related genomes, and an outgroup. (a) the tree, with branch lengths given by the genetic distance (the estimated number of mutations). The uploaded sequence is indicated by a red dot. (b) the collection date of the sequence within each tip group, coloured by location. (c) The frequency of occurrence of spike mutations within each tip group. (d) The collection date of the sequence within each tip group, coloured by lineage (data analysed by GISAID, Audaciy Instant).

## Discussion

The identification of common mutations such as C3037T, C14408T, and A23403G in the Sylhet region highlights the genetic diversity of SARS-CoV-2 and its evolutionary dynamics. These mutations, which are also prevalent in European strains, suggest potential transmission routes and highlight the importance of global surveillance (Lauring & Hodcroft, 2021).

In our study, we looked at how common and linked five major spike protein mutations found in SARS-CoV-2 were to 37 COVID-19 patients. These mutations are D614G, A570D, D1118H, A222V, and P681R. These mutations showed different prevalences and clinical effects in the variables studied.

### D614G

The spike D614G mutation had the highest prevalence at 94.6% (35/37), consistent with its global dominance (Akter et al. 2023), and is found in multiple variants, including the Alpha (B.1.1.7) and the Delta variants (B.1.617.2) (https://doi.org/10.1038/s41586-020-2895-3). It’s interesting that, even though previous research had suggested that D614G made viruses more contagious, we didn’t find any significant link between it and clinical outcomes like symptoms or hospitalization rate (all p-values > 0.05). However, this high prevalence limits the statistical power to detect potential associations and emphasizes the need for larger, more diverse cohorts in future studies. *A570D and D1118H*

The spike mutations A570D and D1118H had a moderate prevalence of 32.4% (12/37). All of these mutations, which are in the S2 domain and are often linked to the Alpha (B.1.1.7) variant (https://doi.org/10.1038/s41467-022-32262-8), did not have any statistically significant relationships with clinical or demographic variables (all p-values > 0.05). There aren’t any clinical links for A570D, which is especially interesting because many studies have found that the Alpha variant is much more likely to spread, with estimates ranging from 50 to 100% higher than previous SARS-CoV-2 strains. The shorter generation time and higher viral load provide additional evidence of accelerated transmission dynamics (https://doi.org/10.1038/s41467-021-27202-x, doi: 10.1186/s12879-022-07623-9). This implies that the genetic context within the variant may modulate the individual impact of A570D.

### A222V

The spike A222V mutation, present in 29.7% (11/37) of cases, also showed no significant associations with demographic or clinical variables. However, a non-significant trend (p = 0.229) indicated a reduced frequency in patients experiencing diarrhea. We have observed this mutation in several variants, such as the Alpha variant and the 20E/EU1 variant, which are associated with the spread in Europe. It has also been detected in the Delta variant (https://doi.org/10.1371/journal.ppat.1010631, https://doi.org/10.1038/s41467-020-19808-4). Despite the literature reporting gastrointestinal symptoms, including diarrhea, in COVID-19 patients, the precise correlation with the A222V mutation remains poorly documented. Research indicates that approximately 2-10% of COVID-19 patients experience gastrointestinal symptoms, often attributing these symptoms to the overall effects of the virus instead of specific mutations (https://doi.org/10.1007/s40200-022-01002-6). New research suggests that A222V may slightly improve ACE2 binding because it changes the structure of the spike protein by adding valine at position 222. This makes the spike protein more hydrophobic, especially near the receptor-binding domain (RBD). This change could increase the flexibility of the RBD, making it more accessible for binding to ACE2 (doi: 10.3390/v15040856). However, our findings suggest that this does not necessarily lead to significant changes in clinical presentation or disease severity.

### P681R

Perhaps the most interesting results came from the analysis of the spike mutation P681R, which occurred in 13.5% (5/37) of cases. Despite its relatively low prevalence, P681R showed several notable trends. A potential association with gender was observed (p=0.057), with a higher prevalence in females (28.6%) than males (4.3%). Although this mutation is an important determinant of viral fitness for variants such as Delta (doi: 10.1101/2021.08.12.456173), there is no clear evidence in the current literature that there is a direct association between the spike P681R mutation and gender. A higher prevalence was also seen in patients with diarrhea (p = 0.081) and anosmia (p = 0.136), but studies show that these symptoms are not caused by specific mutations(https://doi.org/10.1007/s40200-022-01002-6, https://doi.org/10.3389/fmicb.2022.1027271). P681R’s strategic location at the furin cleavage site is critical for increased viral replication of the delta variant of SARS-CoV-2. This mutation facilitates more efficient cleavage of the spike protein, which is essential for effective virus entry into host cells, contributing to this variant’s increased transmissibility and pathogenicity (10.1016/j.celrep.2022.110829). This makes these trends particularly intriguing for further studies.

## Implications of Viral Mutations and Future Research Directions

The varying prevalence of these mutations in our sample provides a snapshot of viral population dynamics during the sampling period. The high prevalence of D614G (94.6%) contrasts with the lower prevalence of P681R (13.5%), suggesting that the Delta variant was not yet dominant in our study population. This emphasizes the importance of continuous genomic surveillance to track the evolution and spread of SARS-CoV-2 variants.

Our findings indicated that the most frequent nucleotide changes in the Sylhet region were A23403G, C3037T and C14408T. These mutations are part of a haplotype that is widespread in European strains of SARS-CoV-2, indicating a potential link between the virus variants circulating in Sylhet and those prevalent in Europe. This genetic link underscores the critical importance of understanding regional genetic variations of SARS-CoV-2. These Europe-associated mutations in Sylhet highlight the pandemic’s global interconnectedness and the potential impact of international travel on viral spread. This emphasizes the need for robust and adaptable international travel policies to contain the cross-border transmission of new variants (Chowdhury et al., 2020). Furthermore, these findings accentuate the indispensable role of local genomic surveillance programs. Such initiatives are critical for real-time viral evolution monitoring, tracking the emergence and spread of new variants, and detecting the influx of foreign strains. This information is important for targeting public health interventions, developing vaccine development strategies, and assessing the potential impact on the accuracy of diagnostic tests.

This study is limited by the relatively small sample size and the focus on a single region. Although the Bangladeshi context provided valuable insights, future studies could enhance the generalizability of the results by expanding the sample size and geographic coverage. Longitudinal studies that track the evolution of the virus over time would complement the current findings and provide a more dynamic perspective on the progression of the pandemic. In addition, whole genome sequencing could reveal possible mutations beyond the spike protein, providing a more comprehensive genetic profile. These expanded research efforts would contribute to a more nuanced understanding of the behavior of the virus and its impact on public health.

## Conclusion

This study provides valuable insights into the complex interplay between SARS-CoV- 2 spike protein mutations, clinical outcomes and patient demographics. Our analysis of the D614G, A570D, D1118H, A222V and P681R mutations revealed that while these frequently occur together, their direct impact on symptoms is minimal. This suggests a more complicated relationship between viral genetics and clinical manifestations than previously thought. Our findings show that age is a crucial factor for recovery from COVID-19, with older patients showing a slower course of recovery. Remarkably, in our statistical models, we observed gender differences in predictors of outcome: clinical symptoms such as diarrhea and viral genetic mutations were more influential for men, while age and disease progression were more significant for women. These findings emphasize the importance of personalized approaches in the treatment of COVID-19, as well as the need for further research aimed at investigating the correlation between these descriptors and COVID-19.

Compared to anosmia, our study population showed a higher prevalence of male patients and a relatively low incidence of diarrhea. While most spike protein mutations showed no significant correlations with clinical outcomes, the trends observed for P681R, particularly the potential association with gender and certain symptoms, warrant further investigation. The identification of mutations that are also present in European strains (A23403G, C3037T, and C14408T) in the Sylhet region highlights the pandemic’s global interconnectedness.

Our findings highlight two critical aspects for public health strategies: the need for enhanced genomic surveillance to detect emerging variants and the importance of personalized, sex-specific approaches to COVID-19 management. The identification of common mutations in Europe underscores the need for vigilant surveillance, while the observed gender differences in outcome predictions emphasize the value of tailored strategies for patient care.

In conclusion, our research contributes to the understanding of the genetics, clinical manifestations and epidemiology of SARS-CoV-2. The complex relationships uncovered between viral mutations, patient demographic characteristics and clinical outcomes underline the need for a multi-faceted approach to COVID-19. As the pandemic progresses, ongoing research, vigilant genomic surveillance, and adaptive public health strategies remain critical to our global efforts to combat SARS-CoV-2 and mitigate its impact on human health. Future studies with a larger sample size and ongoing genomic surveillance may help clarify the clinical significance of these mutations.

## Author Contributions

SA, ZZ, AH and MAO resources, analyses, writing, and methodology; MS, JINO, GVRS, SA, and AH: investigation, software, validation; TAB, MI, MHS, BG, SFC and SK conceptualization, review; SA editing, visualization, and supervision. All authors have carefully reviewed and consented to the final version of the manuscript.

## Data Availability Statement

The viral genome sequences generated in this study have been deposited in the GISAID database. The accession numbers are as follows: EPI_ISL_1538415, EPI_ISL_1538416, EPI_ISL_1538417, EPI_ISL_1538432, EPI_ISL_1541553, EPI_ISL_1542073, EPI_ISL_1542918, EPI_ISL_1545269, EPI_ISL_1545270, EPI_ISL_1546389, EPI_ISL_1547360, EPI_ISL_1547371, EPI_ISL_1547372, EPI_ISL_1547828, EPI_ISL_1548044, EPI_ISL_1548059, EPI_ISL_1548072, EPI_ISL_1549238, EPI_ISL_1550055, EPI_ISL_2002483, EPI_ISL_1790058, EPI_ISL_1790062, EPI_ISL_1790209, EPI_ISL_1790211, EPI_ISL_1790213, EPI_ISL_1805572, EPI_ISL_1805604, EPI_ISL_1805631, EPI_ISL_1805650, EPI_ISL_1805661, EPI_ISL_1805696, EPI_ISL_1550506, EPI_ISL_1550911, EPI_ISL_1563631, EPI_ISL_1563632, EPI_ISL_1563633, EPI_ISL_1563634. Additional data related to this study, including processed sequence alignments and variant calling results, are available upon reasonable request from the corresponding author.

## Acknowledgments

This work was supported by the Ministry of Science and Technology, People’s Republic of Bangladesh, and Bangladesh Council of Scientific and Industrial Research (BCSIR), Dhaka-1205, Bangladesh.

## Ethics Statement

The protocol for sample collection from COVID-19, Recovered and Healthy humans, sample processing, transport, and RNA extraction was approved by the National Institute of Laboratory Medicine and Referral Center of Bangladesh. The study participants received a written informed consent letter consistent with the experiment. This study was approved by the human research ethics committee of NILMRC (project code: 224125200).

